# Not digested: algal glycans move carbon dioxide into the deep-sea

**DOI:** 10.1101/2022.03.04.483023

**Authors:** Silvia Vidal-Melgosa, Matija Lagator, Andreas Sichert, Taylor Priest, Jürgen Pätzold, Jan-Hendrik Hehemann

**Author notes:** S.V.-M and M.L. contributed equally to this work. Department of Chemistry, Manchester Institute of Biotechnology, University of Manchester, Manchester, M1 7DN, UK. Institute of Molecular Systems Biology, ETH Zurich, 8093 Zurich, Switzerland.

## Abstract

Marine algae annually synthesize gigatons of glycans from carbon dioxide, exporting it within sinking particles into the deep-sea and underlying sea floor, unless those glycans are digested before by bacteria. Identifying algal glycans in the ocean remains challenging with the molecular resolution of conventional analytic techniques. Whether algal glycans are digested by heterotrophic bacteria during downward transport, before they can transfer carbon dioxide from the ocean surface into the deep-sea or the sea floor, remains unknown. In the Red Sea Shaban Deep, where at 1500 m water depth a brine basin acts as a natural sediment trap, we found its high salt and low oxygen concentration accumulated and preserved exported algal glycans for the past 2500 years. By using monoclonal antibodies specific for glycan structures, we detected fucose-containing sulfated polysaccharide, β-glucan, β-mannan and arabinogalactan glycans, synthesized by diatoms, coccolithophores, dinoflagellates and other algae living in the sunlit ocean. Their presence in deep-sea sediment demonstrates these algal glycans were not digested by bacteria. Instead they moved carbon dioxide from the surface ocean into the deep-sea, where it will be locked away from the atmosphere at least for the next 1000 years. Considering their global synthesis, quantity and stability against degradation during transport through the water column, algal glycans are agents for carbon sequestration.

**Significance statement:** Algae and plants use the greenhouse gas carbon dioxide to synthesize polymeric carbohydrates, or glycans, for energy storage, structural support and as protection against invasion by microbes. Glycans provide protection, are carbon sinks and enable carbon sequestration for as long as they are not digested by bacteria or other organisms, which releases the carbon dioxide back in to the atmosphere. In this study, we show that non-digested algal glycans sink into the deep ocean and into marine sediment. Thus, glycans are more than food for animals and prebiotics for bacteria, they are also molecules that remove carbon dioxide from the atmosphere and transfer it to the deep-sea, where it can be stored for 1000 years and longer.

## Introduction

Marine algae synthesize polysaccharides with diverse structures resulting in biological functions that shape their contribution to carbon cycling and carbon sequestration. Algae annually convert gigatons of carbon dioxide via photosynthesis into glucose^1^. Algal enzymatic pathways isomerize the glucose into various carbohydrate monomers. These monomers are polymerized into structurally diverse glycans^2^. Depending on algal species, life stage and nutrient status they contribute up to 80% of the algal organic biomass^3^ and have various biological roles that affect their ability to sequester carbon. Structurally simple, intracellular polysaccharides, such as laminarin and glycogen made of glucose, provide energy and transient carbon storage for days to months^4^. They evolved to be easily synthesized and digested to store and release carbon energy rapidly within hours, visible in their diurnal turnover within diatom cells^5^. Structurally more complex polysaccharides such as alginate, cellulose and mannans make gels or fibrils that surround and form the cell wall^6^, which needs to remain stable at least for the life time of the algae. Structurally among the most complex in terms of building blocks, linkage types and configuration are the secreted cell surface glycans such as fucose-containing sulfated polysaccharide (FCSP) from diatoms and brown macroalgae^7–9^. One of the roles of secreted anionic polysaccharides is to provide a form of defense against bacteria, viruses and potentially grazing animals^10–12^. Because these secreted polysaccharides are - outside of the cell - in direct contact with bacteria, their structure must be not labile and instead be challenging to degrade and metabolize with enzymes, or else bacteria can use this carbon energy to outcompete the algae^13^. This protective function leads to slower degradation rates, enabling their accumulation and carbon storage for months during algal blooms^14^. Whether algal glycans are stable enough to export carbon in sinking particles through thousands of meters of seawater with glycan-digesting bacteria, remains still to be tested.

Together with extensive glycan synthesis by algae comes extensive degradation of glycans by marine bacteria and other organisms, questioning their ability to export carbon dioxide. The biological carbon pump is the gravitational settling of photosynthetic derived organic matter within sinking particles, contributing to the ability of the ocean to regulate climate. About 1-5% of the organic matter that sinks out of the surface ocean reaches the sea floor, indicating extensive degradation takes place during transit in the water column^15^. Assuming non-selective degradation, as observed in sinking particles^16^, glycans would be consumed at a rate proportional to other molecules present in algae-derived organic matter. These low molecular resolution data suggest that between 95-99% of the glycans synthesized by algae in the sunlit ocean and exported by the biological carbon pump are degraded by heterotrophic processes before they can reach the ocean floor. Intense degradation is consistent with proteomics and metagenomics of bacterioplankton during algal blooms showing marine bacteria synthesize glycan-degrading enzymes, including glycoside hydrolases, polysaccharide lyases, carbohydrate esterases and sulfatases^17,18^. Incubation studies of glycans added to seawater from different oceanic basins, depths and sediments confirm degradation by bacterial enzymes, at glycan type-specific rates^19–21^ where the secreted, structurally more complex FCSP is degraded slower than the structurally simple β-glucan laminarin. Algae-derived organic matter in sinking particles experiences intense remineralization from zooplankton grazing and from particle-associated microbes in the water column, potentially limiting export of glyco-carbon^16,22,23^. Furthermore, once carbohydrates reach the sediment surface, active benthic organisms and microbial communities continue to degrade organic matter by both aerobic and anaerobic remineralization^24,25^. Despite degradation, once carbon-rich particles reach sufficient depth (∼1000 m) and depending on location, the released CO2 takes up to 1000 years to reach back to the surface with the ocean overturning circulation^26,27^. Before we are able to accurately predict or can influence the size of the biological carbon pump to sequester carbon, we need to know which types of organic molecules resist degradation. In this context, it is important to know if algal glycans are stable enough against glycan digesting bacteria to reach a water depth where carbon can be stored for 1000 years or longer.

Bioanalytic approaches can resolve the structures of glycans in marine dissolved and particulate organic matter and shed light on their contribution to carbon sequestration. We have recently identified several polysaccharide types, including FCSP, β-glucan, β-mannan and arabinogalactan, during microalgal blooms in the North Sea. The FCSP synthesized by diatoms was stable for weeks to month and formed particles^14^. Here, we test the hypothesis that FCSP and other algal glycans export carbon dioxide within sinking particles into the deep-sea. We investigated sediment from a deep-sea anoxic brine in the Red Sea, where high salinity and absence of oxygen^28^ preserves organic molecules against biological degradation. To identify individual glycan structures, we used carbohydrate microarrays combined with glycan-specific monoclonal antibodies (mAbs). We detected ten glycan epitopes, some of which are exclusive to algae. Their presence in sediment indicates intact algal glycans export carbon in sinking particles to the deep-sea.

## Results

### General characteristics of the cores and sampling sites

The Shaban Deep is an anoxic brine-filled basin in the Red Sea that enables algal glycan preservation. The Red Sea is mainly surrounded by desert and semi-desert land without major rivers introducing glycans from terrestrial plants. Its sediments are mainly formed from dust and deposits of four phytoplankton groups: diatoms (present throughout the year and especially abundant in winter), dinoflagellates (dominate during fall), nanoplankton and blue-green algae (mostly present in late spring and summer)^29^. The Shaban Deep brine basin has a salinity of ∼260 ‰, a temperature of ∼24 ºC and is depleted in dissolved oxygen (<0.3 mg L^-1^), which excludes the alteration of organic matter by benthic macrofauna^28,30^. Together, these characteristics increase the chance to detect non degraded, algal glycan structures. We hypothesized that the core GeoB7802-1 (named anoxic), which was sampled from within the anoxic brine-filled basin (**Fig. S1 and Table 1**), would preserve the algal glycans.

**Table 1.**
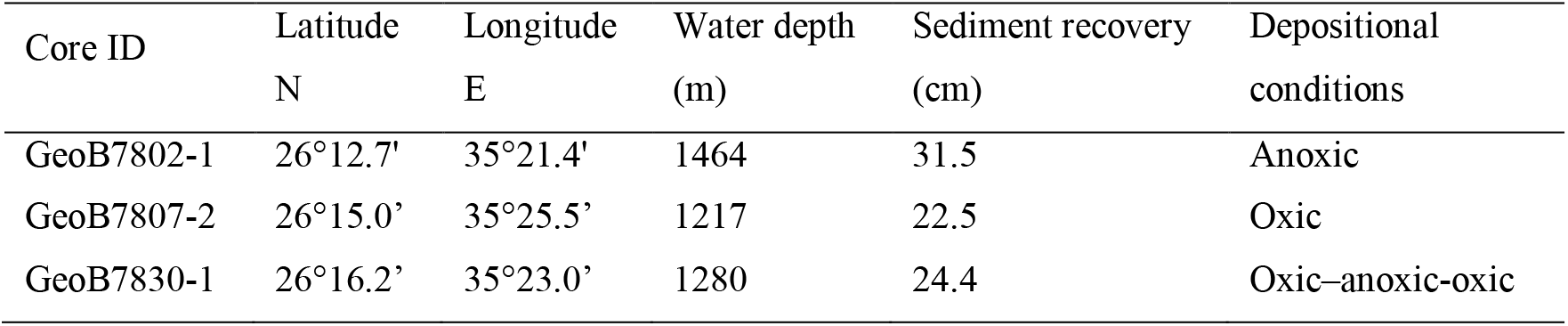
Core sampling sites in the northern Red Sea and water depths in these locations. Cores were collected with the R/V Meteor in the cruise M 52/3, March 2002^30^. Depositional conditions are based on the cores location as well as presence of lamination.

| Core ID | Latitude<br>N | Longitude<br>E | Water depth<br>(m) | Sediment recovery<br>(cm) | Depositional<br>conditions |
| --- | --- | --- | --- | --- | --- |
| GeoB7802-1 | 26°12.7' | 35°21.4' | 1464 | 31.5 | Anoxic |
| GeoB7807-2 | 26°15.0' | 35°25.5' | 1217 | 22.5 | Oxic |
| GeoB7830-1 | 26°16.2' | 35°23.0' | 1280 | 24.4 | Oxic-anoxic-oxic |

For reference to core GeoB7802-1 (anoxic), we selected core GeoB7807-2 (named oxic) and core GeoB7830-1 (named oxic-anoxic-oxic, OAO), which were both obtained in close proximity to the brine (**Fig. S1 and Table 1**). The oxic core was sampled from an oxic environment, where burrowing benthic macrofauna rework the upper cm of the sediment, mix its components, influence the microbial community structure and typically enhance organic matter degradation rates^31,32^. The oxic-anoxic-oxic core was chosen because it was formed under changing conditions, with the Shaban Deep brine surface level increasing and decreasing^33,34^ the core experienced oxic and anoxic depositional conditions. At the time of sampling, this site was located above the brine-seawater interface^30^. For later discussions, we define three “fractions” based on the sediment color and presence of lamination (**Fig. S2**). The upper oxic fraction from surface to 7 cm (mainly light brown), the central anoxic fraction from 8 to 19 cm (laminated, with black to dark grey layers and light layers) and the lower oxic fraction from 20 to 24.4 cm depth (orange to light grey).

The cores were sectioned into a total of 105 sediment sample layers: 33 from the anoxic, 35 from the oxic and 37 from the OAO core (see sectioning criteria as well as further characteristics of the cores, brine-seawater interface depth and age of the cores top and bottom layers at SI Appendix, Supplementary Text and **Table S1**).

### Anoxic brine preserves organic carbon

The anoxic brine environment preserves organic carbon. We analyzed the total organic carbon (TOC) content of all three sites and found that the anoxic brine contained the highest amount. While most of the anoxic core layers were dominated by dark, diatom-rich material^33,35^, the layer at 28 cm depth contained primarily light, coccolith-rich material (**Table S2**). This broad light laminae layer had the lowest TOC value (0.33%) as well as the highest calcium carbonate content (60.31%, **Table S3**), which is a main component of coccolithophore shells (coccoliths)^36^.

Although TOC wt.% (percentage TOC per sediment dry weight) in the anoxic core varied between layers, there was no decrease with depth and average values throughout the core were 1.74 ± 0.37% (**Fig. 1A**). These values are comparable to the central anoxic fraction of the OAO core with 1.76 ± 0.46%. In contrast, the average TOC values of the upper (0.24 ± 0.14%) and the lower (0.12 ± 0.03%) oxic fraction of the OAO core were similar to the values of the uppermost 5 cm oxic core with 0.21 ± 0.07% (**Fig. 1B,C**) and higher than the bottom cm (6 to 22.5 cm depth) of the oxic core, which had values of 0.09 ± 0.01%. In conclusion, TOC content was more than eight times higher for the anoxic core from within the brine than for the oxic core outside of the brine (**Fig. 1A,B**), indicating that the anoxic brine environment preserves organic carbon.

**Fig. 1.**
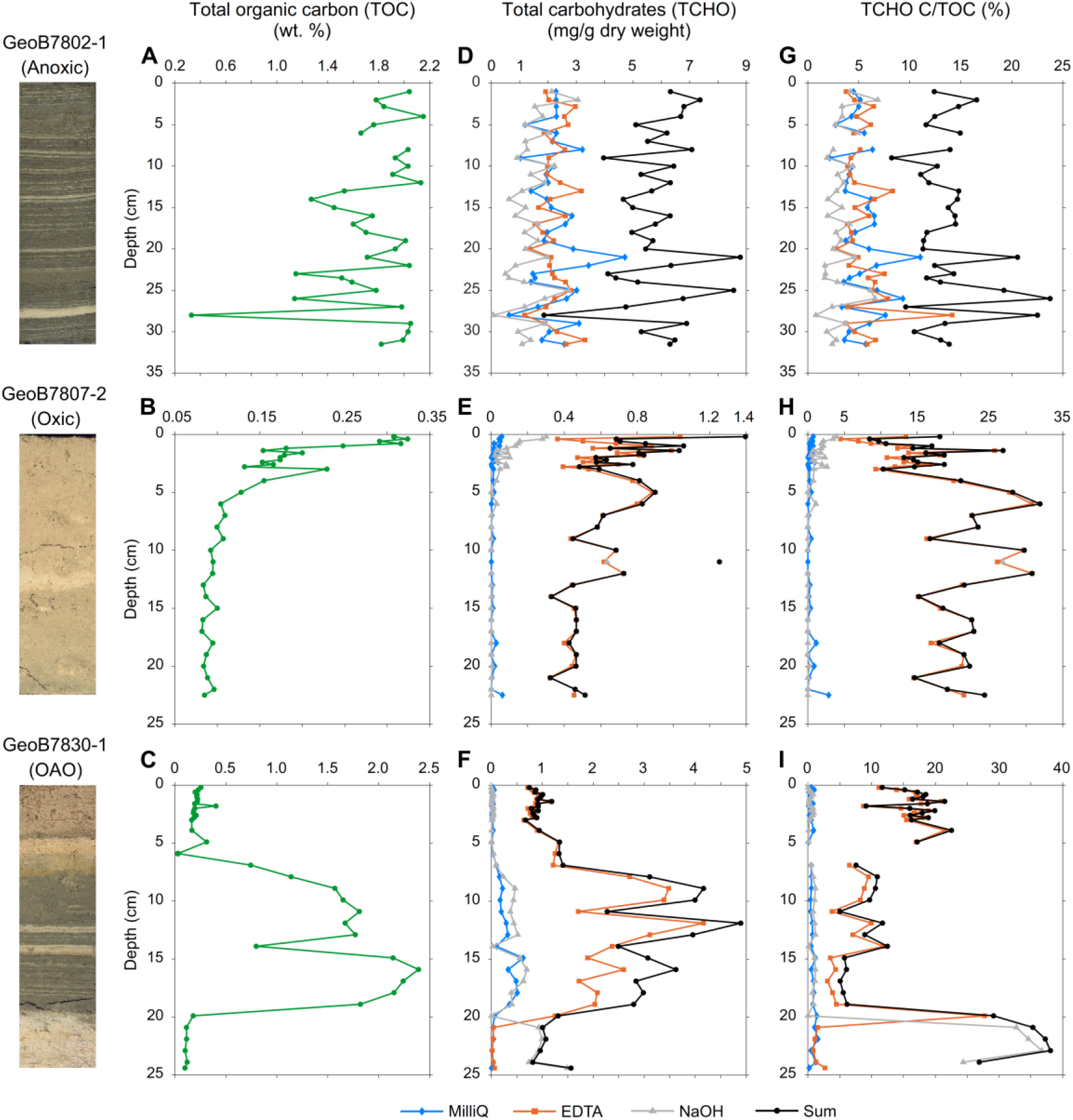
Anoxic brine conditions preserve different carbohydrate types in marine sediment. Depth profiles of total organic carbon (TOC) and total carbohydrates (TCHO) in the three cores. Scans of representative cores (collected with the multicorer at the same time and location as the cores analyzed) are displayed at the left of the corresponding panels. TOC data **(A-C)** are presented as percentage of TOC per sediment dry weight, i.e. TOC per 100 mg dry sediment. **(D-F)** TCHO concentration of all extracts was measured (see color legend) and “Sum” corresponds to the addition of them per layer in the anoxic **(D)**, oxic **(E)** and OAO **(F)** cores. TCHO is expressed as mg carbohydrate as glucose equivalent per g sediment dry weight. **(G-I)** The ratio of TCHO carbon to TOC, i.e. mg of carbon from carbohydrates to mg TOC in dry sediment. In panels (E and H) the NaOH value at 11 cm depth is most likely an error and is shown without connecting lines to the neighboring layers (for panel H sum value at this depth was 52.76). OAO: oxic-anoxic-oxic. Depth, cm below surface.

### Anoxic brine preserves glycans

We used different solvents to extract glycans based on their physicochemical properties, such as solubility, and to separate them from each other and from metal ions and minerals^37,38^. Among the tested parameters (see **Table S4**), extraction with a sequence of MilliQ, EDTA and NaOH, combined with sonication yielded the highest amount of extractable carbohydrates (see schematic in **Fig. S3**, details in Material and methods). Preservation of the glycan polymeric structures in the anoxic core extracts was later confirmed through detection by glycan-specific mAbs.

Next, we quantified the total carbohydrates (TCHO) in the sediment extracts using the phenol-sulfuric acid method^39^. We found the oxic core contained lower TCHO sum values than the anoxic core (**Table S5**) and additionally our data show glycan composition in the anoxic versus the oxic core differed. While the anoxic core (**Fig. 1D**) contained similar amounts of MilliQ, EDTA and NaOH extractable glycans, in the oxic (**Fig. 1E**) and OAO (**Fig. 1F**) cores the TCHO in MilliQ and NaOH extracts were depleted compared to EDTA. In addition, in the anoxic core the concentrations of carbohydrates were similar throughout the core, with averages of 2.21 ± 0.79 in MilliQ, 2.22 ± 0.49 in EDTA and 1.40 ± 0.64 mg g^-1^ in NaOH (**Fig. 1D**). Continuous presence of carbohydrate in the three solvents indicates that a variety of glycans with different physicochemical properties reached the brine-sediment interface during the last two millennia. In contrast, the EDTA extractable carbohydrates makes up to >98% of the total glycans found in the oxic and OAO cores (**Fig. 1E,F**). Low amounts of MilliQ and NaOH extractable carbohydrates were detected in the upper 1 cm and 3 cm of the oxic core and in the OAO anoxic fraction, while mainly the NaOH ones were found in the lower oxic fraction of OAO core. In the anoxic core, the carbohydrate types extracted with the three solvents were preserved while in the oxic core only a fraction of EDTA extractable carbohydrates resisted degradation.

The total carbohydrate carbon relative to sediment organic carbon (mg of carbon from carbohydrates to mg TOC) average was comparable between anoxic (14 ± 3%), oxic (19 ± 6%) and OAO upper oxic (17 ± 4%) cores (**Fig. 1G-I**). The OAO central anoxic fraction contained 8 ± 3% and the OAO lower oxic fraction 33 ± 5%. It should be noted that the 33% ± 5% number might be inflated by dividing through the much smaller TOC values of this fraction. In these layers we could extract a higher proportion of polysaccharide with NaOH, indicating presence of crystalline or water insoluble polysaccharides. Percentage values found in the anoxic and oxic cores are comparable to values found in other sediment samples of the Pacific, Atlantic, Baltic Sea and other sites^23,40,41^. Note that while TOC data include all molecular sizes, our TCHO data include sizes >1 kDa, thus monosaccharides and short oligosaccharides are not represented.

### Monosaccharides indicate related glycans in sediments across sites

Monosaccharide composition of bulk glycans exposes potential origin and mechanisms of persistence in the sediment, albeit at low molecular resolution. The carbohydrates present in the MilliQ and EDTA extracts of all cores were hydrolyzed with acid into monosaccharides, which were identified and quantified by high performance anionic exchange chromatography with pulsed amperometric detection (HPAEC-PAD) (**Fig. 2**).

**Fig. 2.**
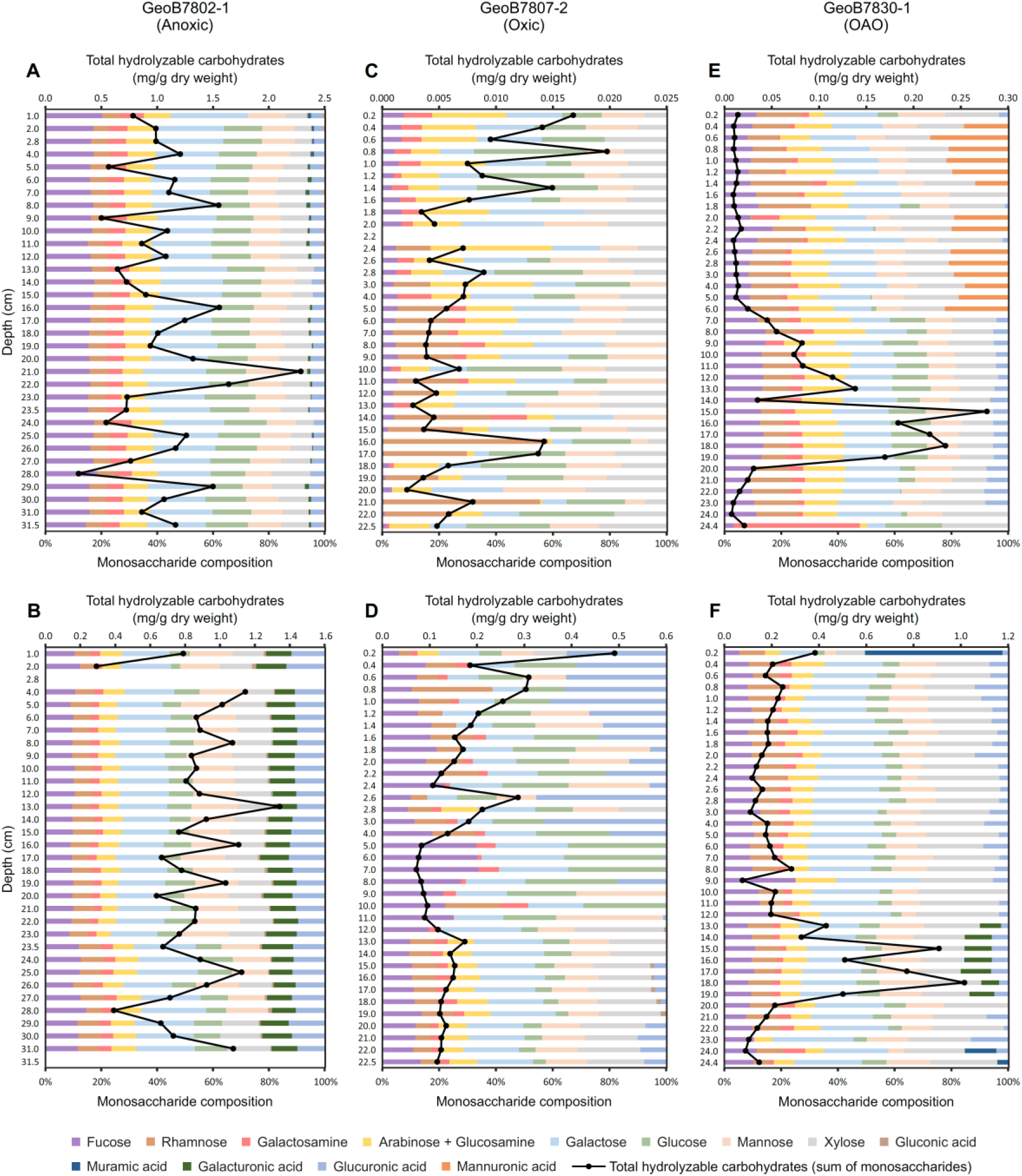
Glycans preserved in each layer of the anoxic brine core share monosaccharide composition. Depth profiles of monosaccharide composition as relative abundance in mol percentage, bottom *x*-axis. Total hydrolyzable carbohydrates (black line) correspond to the addition of all monosaccharides per each layer as mg per g sediment dry weight, top *x*-axis. MilliQ (top) and EDTA (bottom) extracts correspond to the anoxic **(A-B)**, oxic **(C-D)** and oxic-anoxic-oxic, OAO **(E-F)** cores. Depth, cm below surface.

The conserved monosaccharide composition of the anoxic core suggests a constant input of similar glycans for over 2500 years. Using a non-metric multidimensional scaling (NMDS) analysis, with the relative abundance of monosaccharides in each sample as input, we found the anoxic core MilliQ and EDTA extract data formed distinct clusters, indicating the composition of each extract remains unchanged with depth (**Fig. 3**). Both extracts shared several major building blocks, i.e. fucose, galactose, glucose, mannose and xylose, and only galacturonic acid and glucuronic acid were enriched in the EDTA extracts (**Fig. 2A,B**), as further discussed below. Despite varying total hydrolyzable carbohydrate concentrations (which correspond to the addition of all monosaccharides per each layer) among layers, the relative monosaccharide composition remained constant in the anoxic core. This compositional homogeneity indicates that input organisms and potential structural alteration and degradation by microbial digestive enzymes were comparable for over 2500 years.

**Fig. 3.**
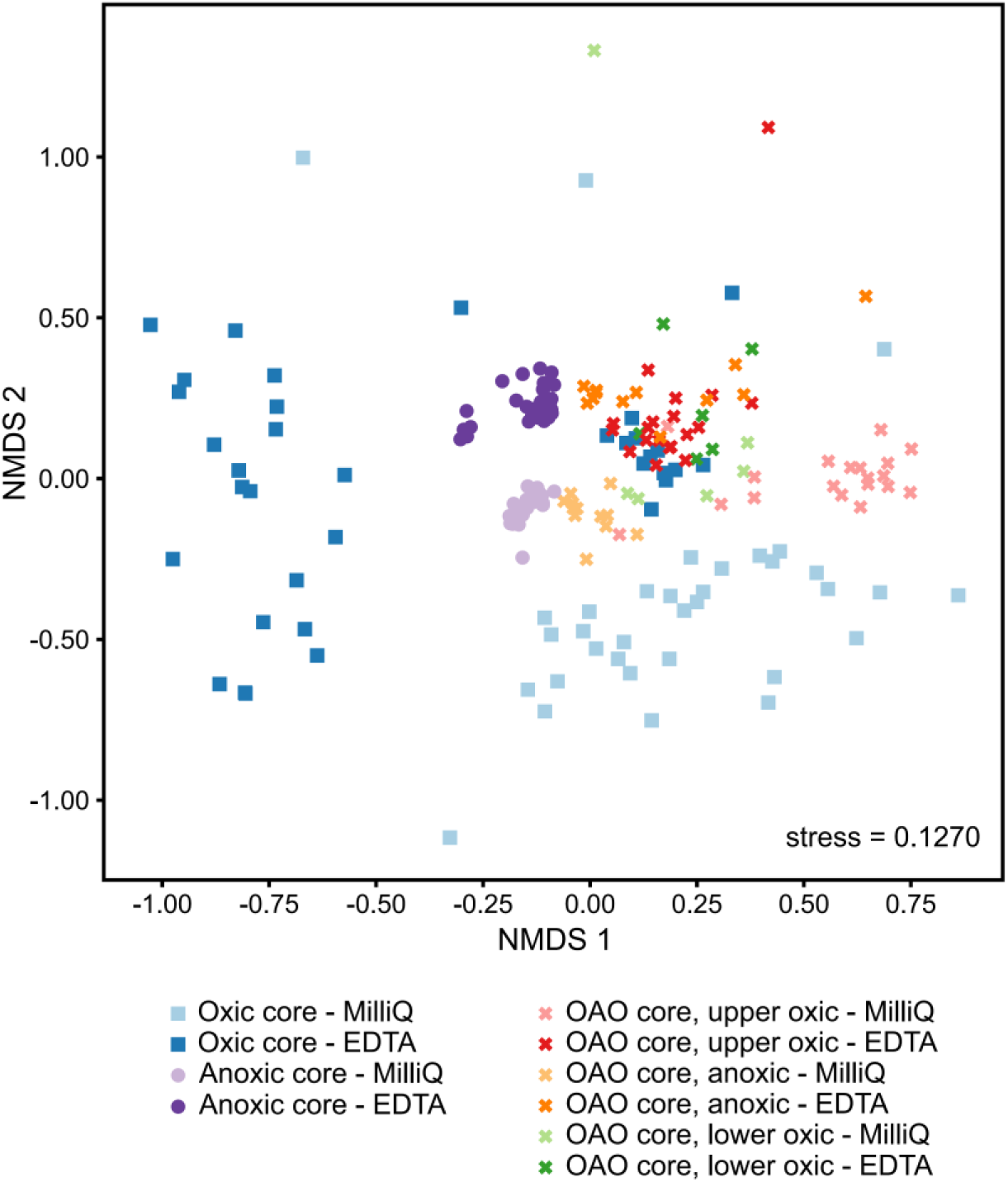
Monosaccharide composition similarities between cores point toward a common glycan source. Non-metric multidimensional scaling (NMDS) plot showing Bray-Curtis dissimilarity of monosaccharide composition in MilliQ and EDTA extracts across the different layers of the three cores. Values were derived from the relative abundance of monosaccharides in 207 sediment extract samples.

This continuity is further evidenced by carbohydrate compositional similarity between the central anoxic fraction of the OAO core and the anoxic core. In the NMDS, all MilliQ extracts of the OAO anoxic layers clustered with the anoxic core. In the anoxic fraction of the OAO core there was an apparent bipartitioning where layers 8 to 12 cm had total hydrolyzable carbohydrate values lower than layers 13 to 19 cm (**Fig. 2E,F**), differing to what was observed by TCHO (**Fig. 2F**) but similar to the trend shown by TOC values (**Fig. 2C**). The EDTA extracts from the OAO anoxic layers with highest total hydrolyzable carbohydrate concentrations, 13 to 19 cm depth (**Fig. 2F**), clustered with the EDTA extracts from the anoxic core (**Fig. 3**). This clustering of anoxic layers across cores further indicates constant input of structurally related glycans that reached the sea floor.

While related glycans appear to have reached the sea floor (**Fig. 3)**, it remains unclear what enabled preservation of EDTA extractable glycans under oxic conditions (**Fig. 1E**). As a chelating agent, EDTA extracts acidic/charged sugars e.g. galacturonic and glucuronic acid bound to cations^37,38^. We compared the amount of acidic monosaccharides in each sediment layer extracted either by EDTA or MilliQ to the total monosaccharide sum of the corresponding extract and confirmed that acidic sugars in EDTA extracts account on average for 14.0% of the EDTA-extracted carbohydrates, 2.6-times more than in MilliQ extracts (*P*-value <0.0001, two-sided *t*-test). Especially in the upper 3 cm of the oxic core, acidic sugars accounted for 20 to 45% of all EDTA-extracted carbohydrates, suggesting interaction with minerals and ions preserves them in the sediment. This preservation is consistent with interactions with metal ions and minerals shielding macromolecules from degradation by rendering them physically less accessible to microbes and their enzymes^42^ or by protecting them chemically because metal ions can activate and also inactivate some enzymes^43,44^ and can be toxic to bacteria^45^.

### Algal glycans move carbon into deep-sea sediment

Although monosaccharide composition can be used to infer the source organism, these associations remain ambiguous, owing to the diversity of glycan structures and their monomer composition. For instance, carbohydrates in sediment dominated by glucose or galactose have been proposed to indicate origin from land plants or macroalgae, respectively^46^, whilst those dominated by fucose were proposed to originate from bacteria^47^. Fucose is a major building block of the diatom and brown macroalgae FCSP and many other non-bacterial glycans. In our anoxic core, glucose, galactose and fucose are dominant, thus without higher molecular resolution their predicted source remains unclear. The monomer composition after acid hydrolysis is an average of all glycans, many of which share the same monomers yet have different structures and origins.

Structure specific mAbs expose algae as a source of glycans in the anoxic brine sediment. To resolve glycan structures with sufficient molecular resolution to identify organisms of origin, we used glycan structure specific mAbs in combination with carbohydrate microarrays. In the anoxic core, the mAbs detected ten glycan epitopes including β-1,3-glucan, β-1,4-mannan, FCSP, xylosyl residues, galactosyl residues on rhamnogalacturonan I, arabinogalactan and alginate (**Fig. 4**). Most of these epitopes have previously been identified during diatom dominated algal blooms in the North Sea^14^. In the oxic and OAO cores, the mAbs did not detect their cognate glycans, not even in the anoxic fraction of the OAO core, despite being related in the NMDS analysis by monosaccharide composition with the anoxic core. The glycans in the oxic and OAO cores underwent degradation or structural alteration by unknown processes rendering the glycan signal non readable by our set of mAbs.

**Fig. 4.**
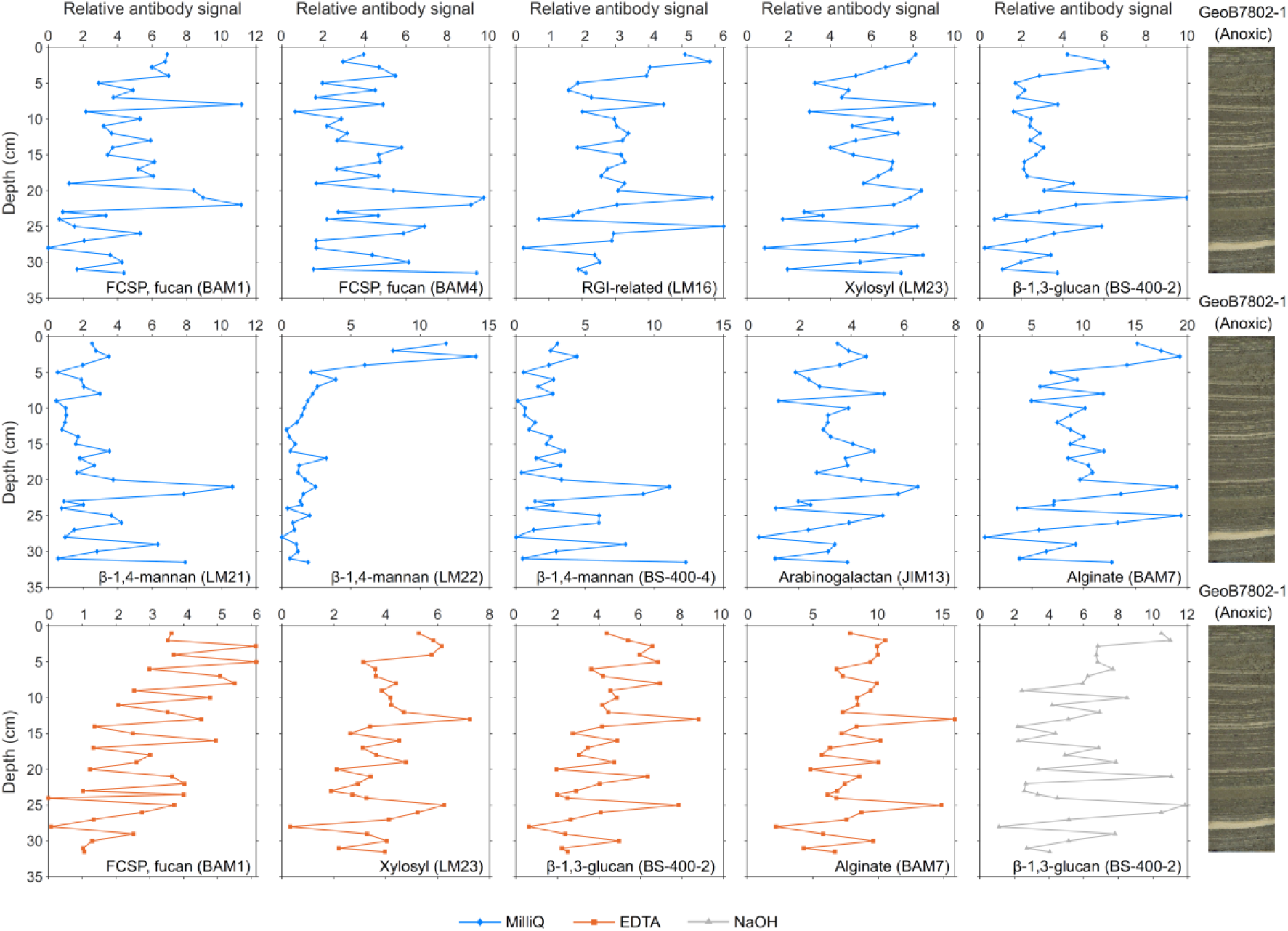
Not digested algal glycans moved carbon dioxide into the deep-sea for the past 2500 years. Shown are polysaccharide structures detected in the anoxic brine sediment (core GeoB7802-1) and their depth distribution. Plots show the relative abundance of polysaccharide epitopes (relative monoclonal antibody (mAb) signal intensity) detected in the different layers of the core. The mAbs (depicted in parenthesis) and the polysaccharide epitope they bind to are shown in each individual plot. A cut-off of 4 arbitrary units was applied and the figure shows all profiles where in at least two depths antibody positive signal (value ≥ 4) was detected. Values lower than 4 are shown for clarity. Details of probes used are provided in Table S6. Depth, cm below surface.

In the anoxic core, glycan structures were present from the top to the bottom. The signal intensity of a single mAb is semi-quantitative but correlates with the relative abundance of its glycan epitope in the sediment layers (**Fig. 4**). Because each mAb has a unique avidity to its recognized epitope, the signal intensities from different mAbs must not be compared to infer abundances between glycan epitopes^37^.

mAbs bound to glycans from the anoxic core, which was previously found to contain diatom-derived matter. All the mAbs had a minimum signal value at 28 cm depth in the coccolith-rich layer, while glycan epitopes found in the MilliQ extracts (except for mAb LM22) had one signal maximum at 21 cm depth that corresponds to a dark diatom-dominated layer. This result is consistent with the mAbs binding to glycans with diatom origin. In general, minima, maxima and overall intensity abundance trends differed between epitopes. An example of different trends is the binding of the three anti-β-1,4-mannan mAbs to the MilliQ extracts. Two of the epitopes (recognized by mAbs LM21 and BS-400-4) presented higher abundances in lower depths, while the distinct epitope targeted by mAb LM22 was mainly present in the upper 4 cm of the core.

Most of the different glycan structures were detected in the MilliQ extracts, whereas EDTA extracts contained signals for FCSP, xylosyl residues, β-1,3-glucan and alginate, and the NaOH extracts for β-1,3-glucan. Glycan epitopes extracted with EDTA and NaOH were also present in the MilliQ extracts, where their abundances were higher -except for β-1,3-glucan that had highest abundance in NaOH. The detection of the same epitope in multiple solvents can be due to the mAb-recognized epitope being a part of different polysaccharides (each released by a particular solvent) that co-occur in the sediment. Alternatively, this can be a result of the intrinsic heterogeneity of glycans, where one polysaccharide type can have differences in length, degree of branching, degree of sulfation and/or methylation influencing its physicochemical properties. Some of these intrinsic differences can promote intermolecular interactions that are challenging to resolve (compared to single polymers) with solvents. Although some of the glycans interact by physical crosslinks that form macromolecular networks, their shown extractability (**Fig. 4**) makes these polymers accessible to analytic methods with increasing molecular resolution.

## Discussion

We tested if glycans synthesized by algae in the sunlit ocean contribute to carbon dioxide sequestration in the deep-sea. By using carbohydrate-specific mAbs, we identified intact algal glycans in sediment. Structural alteration by microbial enzymes would modify the glycan epitope resulting in no mAb signal. Even with partial degradation to oligosaccharides there would be no signal, as our microarray approach allows immobilization of polysaccharides but not of monosaccharides and short oligosaccharides^48,49^. Thus, the microarray data demonstrate export of intact algal glycans directly to the deep-sea floor. Under oxic, non-brine conditions the native algal glycan signal was not detected, suggesting they were degraded or partially degraded by the active benthic microbial communities.

Glycans are preserved in sediment under high salt and low oxygen concentrations. The three cores investigated in this study were harvested in close vicinity at comparable depths, thus the organic matter that reached the sea floor experienced similar synthesis and degradation processes at the sea surface and during sinking through the water column. The lower TCHO concentrations found in the cores from outside the brine indicate degradation at the sediment surface^24,25^. Importantly, although the TCHO values were lower than those in the anoxic brine core, EDTA-extracted carbohydrates were present throughout the oxic core. Carbohydrates released with the chelating agent EDTA are negatively charged and interconnected by ion crosslinks with calcium and other metal ions^50^ or are associated to these cation-dependent networks. Physical crosslinking between glycan polymers creates supramolecular networks that can make them inaccessible to microbes and their enzymes^51,52^. Also the interaction of these carbohydrates with metal ions could shield and protect them. Both mechanisms have been proposed for the preservation of organic matter in marine sediment^40,42,53^. Another possibility, which, to the best of our knowledge, has not yet been discussed in the context of organic matter preservation but which merits consideration, is that in the laboratory and *in vivo* some metal ions are inhibitors of enzymes, including bacterial glycoside hydrolases^44,54^.

Multiple different glycan epitopes were detected in the anoxic brine sediment. Some of the detected glycans are found in land plants and marine algae, including β-1,3-glucan (callose and laminarin, respectively), β-1,4-mannan, xylosyl residues, galactosyl side chains on rhamnogalacturonan I and arabinogalactan^14,55^, see references associated to land plants in **Table S6**. The sediment, including its glycans, is an assemblage of microalgal matter dominated by diatoms^33,35^. Diatoms and other microalgae as the source is consistent with the detection of structurally related glycans during North Sea diatom blooms^14^. In the case of alginate, the only known sources are brown macroalgae and two genera of terrestrial bacteria. For BAM7 (mAb specific to alginate), cross-reactivity with pectin and, to a lesser extent, fucan was reported^56^. This cross-reactivity raises the possibility that the mAb was binding to a fucan epitope. In conclusion, most of the structures were found throughout the brine core, indicating export by diatoms and other microalgae for the past 2500 years.

Although bacterial enzymes for glycan degradation are abundant in metagenomes of bacterioplankton during algal blooms^17^, it remains unclear to what extent this potential for glycan degradation is used by marine bacteria. Our microarray data reveal several algal glycans reached the deep-sea floor. Genes coding for carbohydrate-active enzymes with putative activity for some of the here identified glycans have been identified in marine *Bacteroidetes*^57,58^, *Gammaproteobacteria*^55,59^ and *Verrucomicrobiae*^60^. Best understood is the degradation pathway for β-1,3-glucan, which was identified in the anoxic brine core (**Fig. 4**). β-1,3-glucans include laminarin, a microalgal storage compound abundant in diatoms and in particulate organic matter in surface water. Laminarin has also been detected in sinking particles, where it contributes to the export of carbon out of the surface ocean^5^. This glycan is labile, with only three enzymes required for its degradation^61^. Bacteria with these enzymes are abundant in the ocean. When present inside microalgae cells, laminarin is protected from degradation by bacterial enzymes. Once it becomes part of the dissolved organic matter pool (for instance due to cell lysis or grazing), it is rapidly degraded by bacteria^62^.

In the surface ocean, diatoms and other phytoplankton thrive and coevolve with bacteria and viruses, which extract energy and nutrients by infecting algal cells and trying to digest them from within and from the outside. FCSP, which covers the surface of diatom cells, resists this degradation^14^. Sulfated fucans from the evolutionarily-related brown macroalgae consist of an α-1,3-or α-1,3;1,4-linked-L-fucose backbone. The backbone is modified with extensive sulfation and side chains with fucose and/or other monomers^9^. Depending on species, physiology and season, algae modify the FCSP resulting in different structures and consequently increased molecular diversity. Out of six tested brown algae, each one had a differently modified FCSP structure^9^. Each one of these FCSPs required bacteria to have an adapted bacterial enzyme degradation system, consisting of dozens of enzymes to extract carbon^60^. Small side chain modifications by acetylation or sulfation make the enzyme sets inactive, resulting in a recalcitrant substrate against non-adapted enzyme sets. This molecular diversity, for which the drivers remain unknown, may provide a molecular lock for carbon sequestration.

Diatoms synthesize FCSP that resists bacterial degradation for weeks to months^14^, theoretically providing sufficient time to reach the deep-sea where carbon dioxide can be locked away from the atmosphere for 1000 years and longer. This hypothesis is here supported by the presence of FCSP in the anoxic brine core. Secreted FCSP forms a gel around diatom cells and also assembles into particles^14^, incorporating and protecting organic molecules against digestive enzymes, akin to how a mucin preserves the organic carbon of a living intestinal cell against invasion and degradation by gut bacteria^11,63^. We consider the carbon preserving function of secreted, cell wall polysaccharides protects labile glycans such as laminarin within algae cells sinking alone or within particles. In addition, the anionic character of FCSP with carboxyl and sulfate groups, which enables gel formation, also enables binding to minerals and ions that can protect the glycans. In conclusion, transported within sinking particles, several algal glycan structures reached the deep-sea, implying an unexplored mechanism enabling preservation of labile and stable glycans in sinking particles and therefore export of carbon dioxide by the biological carbon pump.

## Materials and methods

### Core processing

The three sediment cores analyzed in this study (**Table 1**) were harvested in the vicinity of the Shaban Deep, in the northern Red Sea, during the RV Meteor cruise M52/3 in 2002^30^. The cores were sampled with a Multicorer (MUC, GeoB), with a diameter of 10 cm, and were immediately stored at 4 °C. An intact sediment-bottom water interface and the upper dm of sediment (see sediment recovery for each core in **Table 1**) were acquired. Cores were obtained from the GeoB Core Repository at MARUM (Bremen, Germany), where they were stored at 4º C since recovery. Entire cores were obtained in the case of GeoB7807-2 and GeoB7830-1, while half of the core (cut in two by the longitudinal plane) in the case of GeoB7802-1. The three cores were sectioned (transverse plane) into 1 cm depth intervals using a knife or a thin cutting plate. Except for the first top 3 cm of the GeoB7807-2 and GeoB7830-1, which were sectioned into 0.2 cm sections. A few exceptions were made in the sectioning of core GeoB7802-1 (anoxic core) to preserve some of the homogeneous layers based on appearance, such as the light layer at 28 cm depth (see exceptions in SI Appendix, Table S2). For all cores, each cut layer was individually stored in a Petri dish, sealed with laboratory film (Parafilm) and stored at 4º C until further processing. Before processing, ¼ of the wet volume of each sample (cut layer) was transferred to a 50 mL tube and frozen at - 80º C (kept as archive sample). In order to get rid of the salt, each sample (non-archive) was dialyzed. Note that to homogenize the samples and get rid of clumps, sediment layers were first transferred to a clean beaker and ∼3 mL of MilliQ were added while mixing the material with a spatula. This allowed for a more effective transfer of the sediment to the dialysis bags. Then, each sample was transferred to a 1 kDa dialysis tubing (Spectra/Por®, Spectrum Laboratories) and dialyzed at 4 °C overnight against 2.5 L MilliQ in a beaker with stirring. The complete volume of MilliQ was subsequently replaced twice, after an incubation period of at least 3 h in between replacements. After the third replacement, the desalted sample was transferred to a 50 mL tube. The bag was washed with a small volume of MilliQ in order to get the residual sediment out. Samples were frozen, freeze dried in a lyophilizer and stored at -80 ºC.

### Core dating

Eight sediment depths were chosen for carbon dating, including the uppermost and near bottom layers of the three sediment cores, along with the estimated top and bottom layer of the central anoxic fraction (based on appearance, presence of lamination) of core GeoB7830-1. About 2 g of wet weight of the archive samples (raw sediment) from the selected layers were washed over a 150 μm mesh sieve under running tap-water. Between samples the sieves were cleaned by sonication as well as by a brush under running water. The washed samples were transferred to separate Petri dishes and dried in an oven at 50 °C overnight. Mesh sieves of sizes 300 μm and 600 μm were used to separate the sediment samples into 3 fractions: < 300 μm, 300-600 μm, > 600 μm. The dried sediment samples were dry-sieved and material from each of the three size fractions was collected in different glass vials. The size fraction 300-600 μm was examined under a Zeiss Stereo.V8 binocular microscope under constant magnification/ Wild M3Z Stereozoom (Leica) microscope. Per each of the eight sediment layers, one hundred specimens of the planktonic Foraminifera species *Trilobatus sacculifer* were selected, picked with a thin paintbrush and transferred to micro-slides. When selecting individuals, the amount of (dis)coloration and damage to the shell was minimized. Samples were analyzed by a Mini Carbon Dating System (MICADAS) at the Alfred Wegener Institute for Polar and Marine Research (AWI) in Bremerhaven, Germany^64^.

### Total organic carbon

For determination of the total organic carbon (TOC), between 2 to 4 g of wet weight of the archive samples (raw sediment) from all core layers were weighed out into 25 mL plastic vessels. Samples were frozen, freeze dried and afterwards individually homogenized using a pestle and mortar. Briefly, once homogenized, the samples were decalcified with acid and then TOC was determined by combustion at 1050º C using a Heraeus CHN-O-Rapid elemental analyzer. Data are given in weight percent dry sediment.

### Carbohydrate extraction

For the three sediment cores, the dialyzed and freeze dried samples from each sediment layer were used for carbohydrate extraction. First, samples were homogenized using a pestle and then ∼ 90 mg of sample were weighed out into a 2 mL tube. As polysaccharides have different extractability and solubility, they were sequentially extracted with three different solvents in this order: autoclaved MilliQ water, 400 mM EDTA pH 7.5 and 4 M NaOH with 0.1% w/v NaBH4. For every 10 mg of sample, 200 μL of solvent was added to the tube (sample extractions were normalized by weight). After vortexing, samples were placed in a sonication bath for 1 h and then centrifuged at 4000 x g for 15 min at 20 °C. Carbohydrate extracts (supernatant) were transferred to new 1.5 mL tubes and the pellet was resuspended in the next solvent, using the same extraction procedure described above. For each core, the three extractions for all layers were performed on the same day. Aliquots of the extracts were immediately used for carbohydrate microarray printing and the rest was stored at -20 ºC. Combusted sand was used as negative control and for each core, two sand samples were included and treated the same as the rest of the samples. Values of the sand controls (average of two, which were always among background signal) were subtracted from the samples.

### Total carbohydrate analysis

Total carbohydrates (TCHO) concentration in the three carbohydrate extracts from all sediment layers was determined. Analysis was performed by the phenol-sulfuric acid method^39^. Briefly, 0.2 mL of each carbohydrate extract was transferred to a 2 mL tube. In addition, glucose standards were prepared and treated in parallel with the extracts. A volume of 0.2 mL of 5% phenol was added to each tube followed by (slow) addition of 1 mL of concentrated sulfuric acid. Samples were mixed for 15 seconds and kept at room temperature for 10 min. Afterwards, they were mixed again and placed in a water bath at 30 °C for 20 min. After incubation, absorbance of the samples and standards was measured at 490 nm with a spectrophotometer. The standard curve was used to determine the carbohydrate concentrations of the extracts as glucose equivalents. Standards were prepared in MilliQ, as we previously prepared glucose standards in MilliQ, EDTA and NaOH and tested the effect of using them on individual extracts (from the three solvents) and there was no variance in the concentration obtained. The TCHO of all extracts from a single core were determined in parallel in the same analysis.

### Monosaccharide analysis by HPAEC-PAD

MilliQ and EDTA carbohydrate extracts (300 μL) were acid hydrolyzed at 100 ºC for 24 h with 300 μL of 2 M HCl (7647-01-0, Analar Normapur) in pre-combusted glass vials (400 °C, 4 h). Subsequently, the acid hydrolyzed samples were transferred to 2 mL tubes. Per each sample a 100 μL aliquot was transferred to a 1.5 mL tube and dried for 3 h at 37 °C at 500 rpm in a centrifugal vacuum concentrator. At this step, 1.5 mL tubes with 100 μL of a monosaccharide standard mix (same as described in^60^) were prepared in 1 M HCl and dried in parallel. The dried samples and standards were suspended in 100 μL MilliQ. Note that to avoid high salt concentrations, all EDTA extracts were suspended in 2 mL MilliQ (1:20 dilution). In addition, to avoid too high monosaccharide concentrations in the MilliQ extracts from core GeoB7802-1 (anoxic core), these extracts were suspended in 500 μL (1:5 dilution). The MilliQ extracts from the other two cores were not additionally diluted (i.e. suspended in 100 μL). All samples were transferred to HPLC vials for analysis. A protocol to determine neutral, amino and acidic sugar concentration using high performance anion exchange chromatography coupled with pulsed amperometric detection (HPAEC-PAD) described previously^65^, was applied. Briefly, samples were analyzed using a Dionex ICS-5000^+^ system equipped with a CarboPac PA10 analytical column (2 × 250 mm) and a CarboPac PA10 guard column (2 × 50 mm). The operation procedure was performed as described in^14^. All dilutions, extra dilution factors and acid hydrolysis dilution (acid hydrolyzed samples were 2-fold diluted when suspending), were considered and data show concentrations in the crude extracts.

### Carbohydrate microarray analysis

All samples from one single core were included in a single run that was printed on the same day of extraction, but extracts from different cores were analyzed independently, resulting in three separate microarray analyses. For a single core, 30 μL of each carbohydrate extract were added into wells of 384-microwell plates, where two consecutive two-fold dilutions were done in printing buffer (55.2% glycerol, 44% water, 0.8% Triton X-100). In order to get rid of bubbles, the microwell plates were centrifuged at 3500 x g for 10 min at 15 °C. A microarray robot (Sprint, Arrayjet, Roslin, UK) was used to print the plates content onto nitrocellulose membrane with a pore size of 0.45 μm (Whatman) under controlled conditions of 20 °C and 50% humidity. A printing replicate per each sample was included. Multiple identical arrays were printed, all containing the same sediment carbohydrate extracts. Each microarray was individually incubated with one of 40 mAbs, which specifically bind to one polysaccharide epitope (see list of probes and their specificity in SI Appendix, **Table S6**). Microarray probing was performed as described previously^49^, where binding of mAbs to their recognized epitope on the microarray is detected by using a secondary antibody conjugated to alkaline phosphatase, which in presence of its substrate produces a colored product. Developed arrays were scanned at 2400 dots per inch and binding signal intensity of each mAb to each sample was quantified using the software Array-Pro Analyzer 6.3 (Media Cybernetics). Data analysis was performed as described previously^49^. In short, per each extract the mean mAb signal intensity was calculated and the maximal mean intensity detected in the whole data set was set to 100 and all other values were normalized accordingly. The highest mean intensity detected in the anoxic brine core was obtained by the mAb LM19 (anti-homogalacturonan) binding to a polysaccharide standard (polygalacturonic acid (Megazyme) at 0.25 mg/mL) that was included as positive control. In addition to positive controls, negative controls for the three extraction solvents were as well included in the print showing no unspecific binding and controls for the secondary antibodies showed no unspecific binding to any of the extracts. A cut-off of 4 arbitrary units was applied.

### Non-metric multidimensional scaling

The non-metric multidimensional scaling was carried out using sklearn v0.24.2 and Python v3.7. The monosaccharide data were normalized and pairwise distances were calculated using the Bray-Curtis distance metric.

### Statistical analysis

The amount of acidic sugars in each sediment layer extracted by MilliQ or EDTA was compared to the total monosaccharide sum of the extract. To test for significant changes, we used a standard independent two-sided *t*-test that assumes equal population variances (getting a *P*-value = 3.8 × 10^−10^), which was performed using SciPy v1.7.3 (scipy.stats.ttest_ind) and Python v3.8.

## Data availability

Data reported in this paper will be deposited in the online data archive PANGAEA and a DOI will be provided before publication.

## Acknowledgments

This work was funded by the Deutsche Forschungsgemeinschaft (DFG) through the Emmy Noether Program grant no. HE 7217/1-1, and through the Cluster of Excellence “The Ocean Floor - Earth’s Uncharted Interface” project 390741603 to J.-H.H.; and by the Max Planck Society. M.L., A.S. and T.P. are members of the International Max Planck Research School of Marine Microbiology (MarMic). We thank the GeoB Core Repository at the MARUM - Center for Marine Environmental Sciences, University of Bremen for providing the three studied sediment cores; the Alfred Wegener Institute MICADAS facility in Bremerhaven for carbon dating analysis. We thank Alek Bolte for his help during sample processing and for HPAEC-PAD measurements; Brit Kockisch for TOC quantification; Lukas Jonkers for his assistance and expertise with processing foraminifera samples for carbon dating; Christoph Häggi and Enno Schefuß for advice and valuable discussions.

## Author contributions

S.V.-M. and J.-H.H. designed research; S.V.-M., M.L., A.S. and T.P. performed research and analyzed data; J.P. supported throughout core processing; S.V.-M., M.L., A.S., T.P., J.P. and J.-H.H. discussed the results; and S.V.-M., A.S. and J.-H.H. wrote the paper.

## Declaration of Competing Interest

The authors declare no competing interest.

### Abbreviations

OAO: oxic-anoxic-oxic core
TOC: total organic carbon
TCHO: total carbohydrates
mAbs: monoclonal antibodies

## Supporting information

Supporting information to this manuscript has been included in the submission material.

## Notes

### Competing Interest Statement

The authors have declared no competing interest.

