## Supplementary Information for "Not digested: algal glycans move carbon dioxide into the deep-sea"

**This PDF file includes:**

Supplementary text
Figures S1 to S3
Tables S1 to S6
Supplementary Information references

### Supplementary Text

#### Further detail on the general characteristics of the cores and sampling sites

The sediment cores analyzed in this study were located at the northern Red Sea. Specifically, we analyzed three cores that were previously sampled from within and from close vicinity to an anoxic brine-filled basin, the Shaban Deep (Fig. S1 and Table 1).

Core GeoB7802-1 (named anoxic) was collected from within the anoxic brine-filled basin (Fig. S1), which consists of four sub-basins that together extend over an area of 10 by 6 km (1). The brine basin has a salinity of ~260 ‰, a temperature of ~24 ºC, is depleted in dissolved oxygen (<0.3 mg $L^{-1}$) and sulfide is absent (2, 3). The anoxic core was formed at a water depth of 1464 m under anoxic depositional conditions, which exclude benthic macrofauna and thus bioturbation. The core was sampled in the southern sub-basin and contains alternating light and dark laminae (Fig. S2), with dark color indicative of high organic matter (OM) concentration. Previous studies on laminated sediment from the Shaban Deep showed that the dark layers are primarily composed of diatom frustules, while the light layers are enriched in coccoliths and terrigenous material (4, 5). The basins of the Shaban Deep have the upper brine-seawater interface at a water depth of 1325 m, with a sharp transition zone and minimal exchange by diffusion and convection at the brine interface (2). The two cores harvested outside the brine were at a similar water depth to the Shaban Deep brine-seawater interface, thus OM degradation and alteration during sinking through the water column would be comparable between the three cores (Table 1). Those two core were chosen as controls to enable comparison of carbohydrate preservation in deep-sea sediment under anoxic brine and oxic non-brine conditions.

Core GeoB7807-2 (named oxic) was sampled at a water depth of 1217 m from a location outside of the brine (Fig. S1) and was formed under oxic depositional conditions. Dissolved oxygen concentrations reported at the same latitude and depth are ~3.2 mg $L^{-1}$ (6), similar to oxygen concentrations measured above the brine-seawater interface of the Shaban Deep (2). Salinity and temperature in the deep layers of the Red Sea are (excluding deep-sea brines) constant and rarely vary from ~40.6 ‰ and ~21.5 ºC (6). These values coincide with the values recorded at the oxic core station during sampling (3) as well as with values measured in deep seawater overlying the brine (2).

Compared to the darker color of the anoxic core, the oxic core was light brown to orange and homogenous with no presence of lamination (Fig. S2).

Core GeoB7830-1 harvested at a water depth of 1280 m was formed under changing conditions as during formation, the site underwent periods with oxic, anoxic brine and finally oxic conditions. This was because the Shaban Deep brine surface level rose and later decreased (4, 7), thus currently the site is located above the brine-seawater interface (3). As a result of the different depositional conditions, this core has one laminated section surrounded by two homogenous intervals at the top and bottom (Fig. S2). We name it oxic-anoxic-oxic (OAO) core. Although transition from the brine to the non-brine environment was likely gradual, we define three “fractions” based on the sediment color and presence of lamination (fraction depths are defined at the main text).

**Core sectioning criteria**

Since we predicted lower OM preservation for the oxic cores from outside the brine (oxic as well as upper oxic fraction of the OAO core), high resolution sectioning was performed on those cores for the upper 3 cm, which were sliced into 0.2 cm depth intervals. This was done to be able to detect potential degradation of OM in the uppermost fractions with greater detail. The rest of those two cores was sectioned into 1 cm depth intervals. As we anticipated higher OM preservation in anoxic brine conditions, the anoxic core had a vertical sectioning resolution of 1 cm from top to bottom with few exceptions (see layers description in Table S2). There were numerous light laminae within the anoxic core that were sectioned together with dark laminae, as a result of the 1 cm slicing. However, the thick light laminae at 27-28 cm depth was sectioned with intent to contain exclusively light material. At 2.8-5 cm and 22-24 cm depth there were turbidite layers.

In the central anoxic fraction of the OAO core, all light layers were sectioned together with dark layers, thus for this core we do not have a sample with only light material.

**Age of the top and bottom layers of the cores**

All three cores were located at water depths deeper than 1000 m (Table 1). The age of the bottom layers of the cores varies. As the input of OM reaching the sea floor was most likely similar due to the close location between them, OM degradation was higher in the non-brine environment; thus the accumulation rate was lower and the $^{14}C$ age of the oxic core bottom was older than the anoxic core even though the oxic core was shorter (Table S1). However, the top layers showed the opposite trend where the anoxic surface layer was about 722 years before present (BP) while the surface layers of the oxic and OAO cores were modern and 22 years BP, respectively. This could be a result of the higher sectioning resolution for the two later ones, where the top 0.2 cm layers were analyzed for $^{14}C$ age while 1 cm layer was analyzed for the anoxic core. In addition, this might be due to the high water content of the brine surface sediments of the Shaban Deep (3). As mentioned in the cruise report, undisturbed surface sediment was recovered from the deep brine basin (3) but after storage the core shrank and thus the first cm of the core contains material that originally was at more than 1 cm depth.

**TCHO content in relation to TOC**

The overall abundance trend for the fully anoxic and fully oxic cores did not show large variations when comparing total carbohydrates (TCHO) concentration to TCHO carbon in relation to total organic carbon (TOC). The main difference in the anoxic brine core was the high value at 28 cm depth (Fig. 1G), which exhibits that the low TCHO value in the "light layer" (Fig. 1D) was as well related to a low TOC content (Fig. 1A). In regard to the fully oxic core, the main difference was in the sum value of the upper couple first cm, where from 0.2 to 2.0 cm depth there was an overall decrease in TCHO values (Fig. 1E) but an increase in TCHO/TOC values (Fig. 1H). This increase is caused as a result of the EDTA extracts, as TOC concentration was much higher in these upper layers but the TCHO EDTA values were not that much higher in the uppermost layers, thus their ratio becomes lower.

In contrast to the above stated cores, in the OAO core opposing values were observed for the three fractions comparing TCHO to the TCHO/TOC ones (Fig. 1, panel F vs. I). This illustrates that the higher TCHO concentration in the central anoxic fraction was accompanied by a higher TOC content, as well as that the proportion of organic carbon corresponding to carbohydrates was higher in the two OAO oxic fractions. In the central anoxic fraction the ratio gave higher values on the upper (less old) layers (Fig. 1I), which results from similar values of sum TCHO but lower values of TOC in these upper layers compared to the lower ones. Based on the TOC, and assuming similar composition of the OM reaching the sea floor, it seems there was additional degradation of OM in the upper layers of the anoxic fraction, which is somehow reflected in the TCHO MilliQ and NaOH values (Fig. 1F). Although this is not reflected in the TCHO

EDTA values, which were even higher in the upper layers, it is suggested by the values and composition of the HPAEC analysis (Fig. 2E,F). In addition, while the two oxic fractions had a similar sum TCHO content, the TCHO/TOC differed with higher values in the lower oxic fraction. This is due to the lower TOC values in the lower oxic fraction - TOC % average of 0.24 ± 0.14 and 0.12 ± 0.03 in the upper and lower oxic fractions, respectively. Note that values at 6 cm depth are not shown (Fig. 1I) as the ratio TCHO/TOC gave extreme values (EDTA extract had a value higher than 100%) because of the remarkably low TOC (0.03%) measured at this depth. Plotting these high values would impede to appreciate the trend of the rest of values of the core. For the same reason, in the bottom layer (depth 24.4 cm) the values of NaOH (60.49) and sum (63.37) are not shown in the plot (Fig. 1I) as they are extremely high because of the lower TOC content and higher TCHO content compared to the layers above.

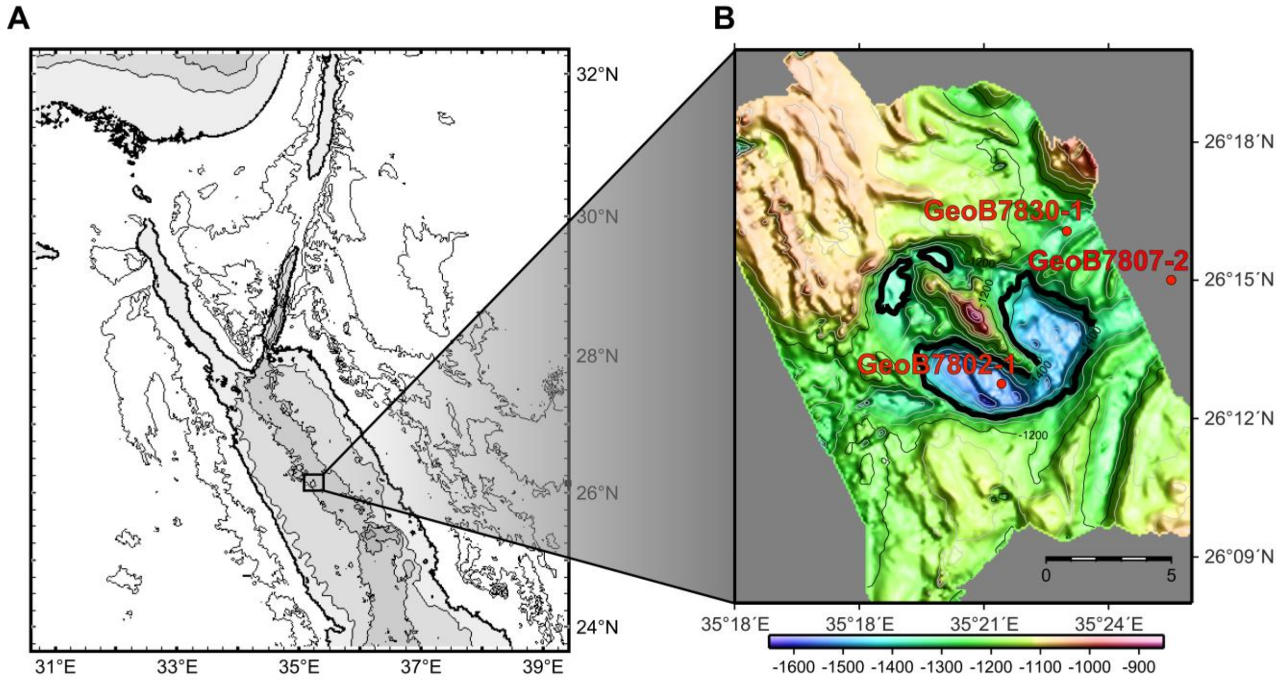


**Fig. S1**. Location of the cores sampling site. (A) Map of the Northern Red Sea where the black square indicates the sampling site region. The map was created with GeoMapApp (www.geomapapp.org). (B) High-resolution bathymetric map of the Shaban Deep (modified after Ehrhardt and Hübscher, 2015 - (8)) displaying the locations of core GeoB7802-1 (anoxic), core GeoB7807-2 (oxic) and Core GeoB7830-1 (oxic-anoxic-oxic, OAO). Black thicker contour indicates the isolines of the Shaban Deep brine interfaces.

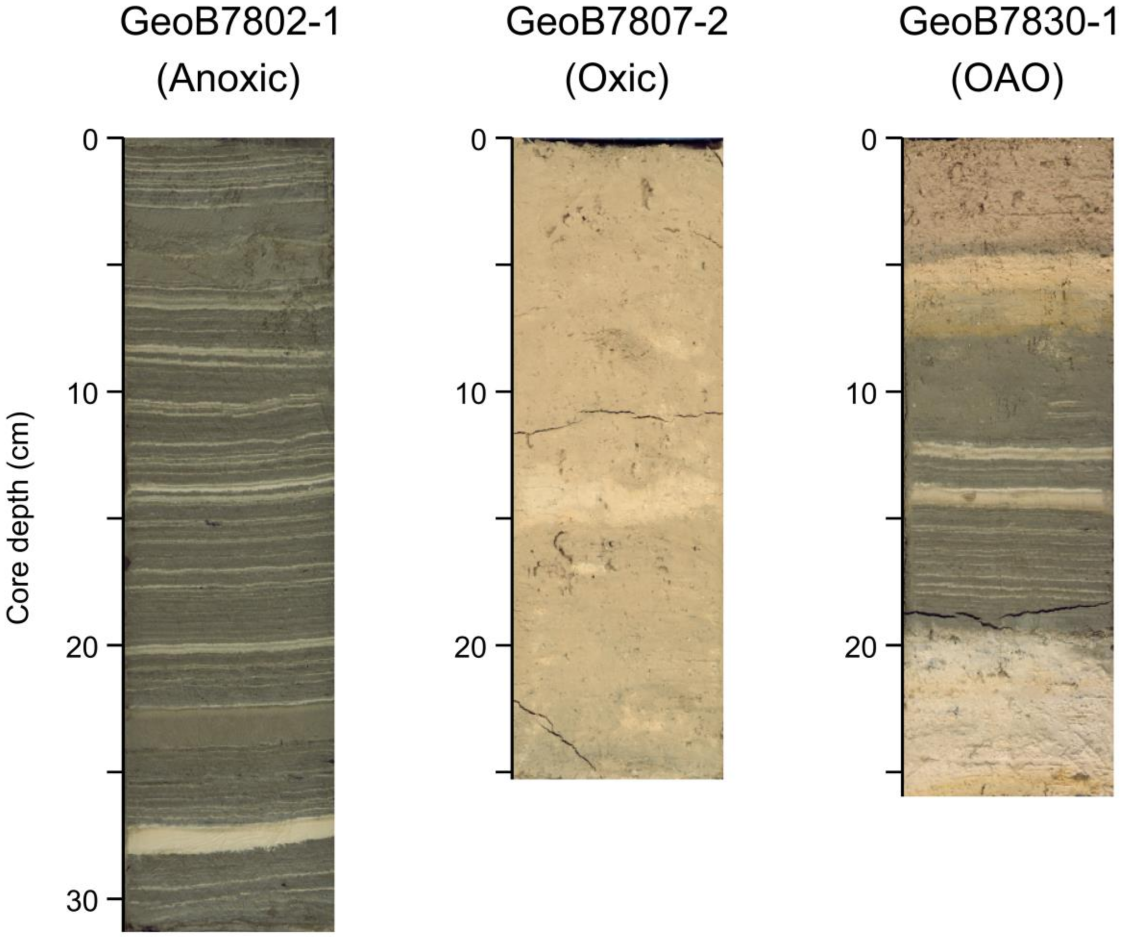


**Fig. S2.** Optical line scans of representative cores. Scans were performed in cores collected with the multicorer at the same time and location as the cores analyzed in this study. These cores were cross-sectioned by the longitudinal plane and scanned with a smartCIS 1600 line scanner system (scans provided by the MARUM GeoB Core Repository). Core depth, cm below surface. OAO: oxic-anoxic-oxic.

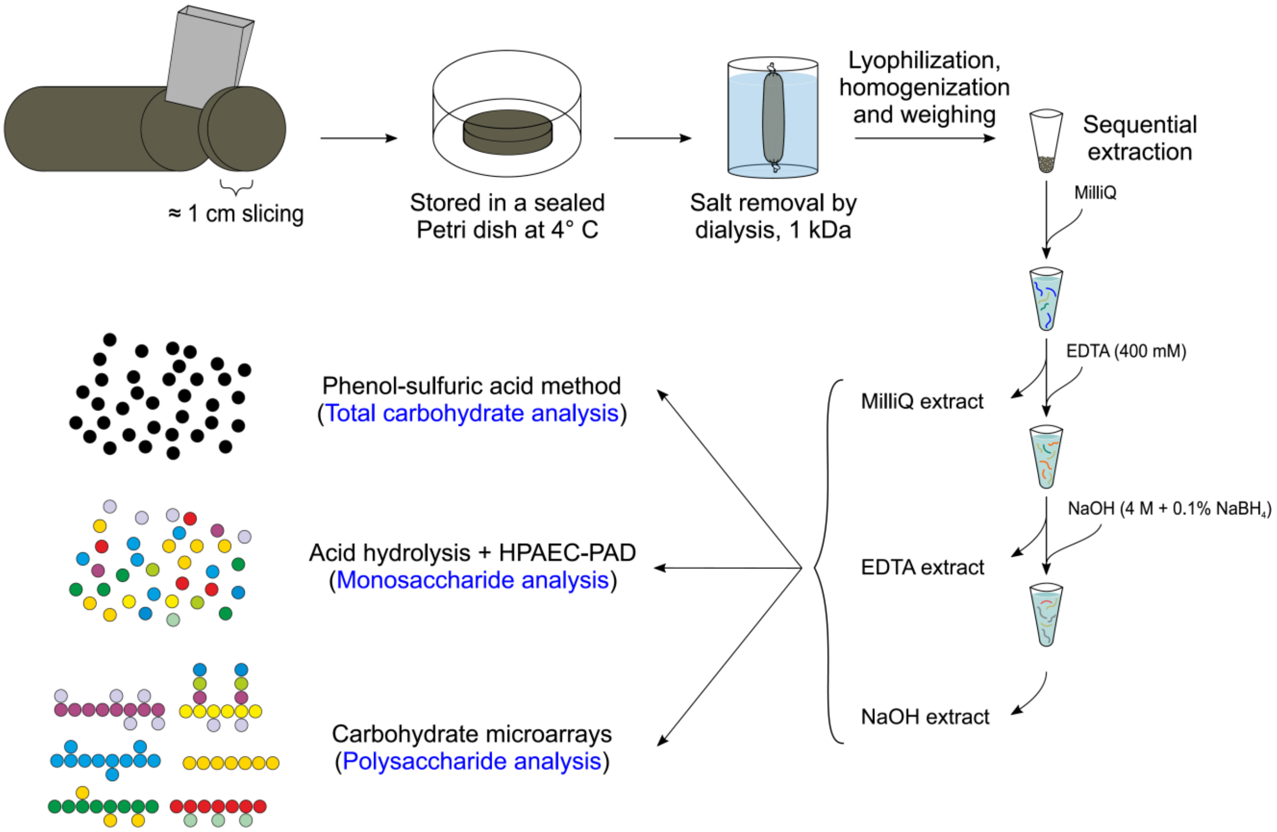


**Fig. S3.** Schematic of the procedure for extraction of polysaccharides developed in this study. Sample processing was followed by what we determined to be the optimal procedure for polysaccharide extraction in anoxic brine sediments, which is sequential extraction by 1 h sonication with MilliQ water, 400 mM EDTA and 4 M NaOH with 0.1% $NaBH_4$. All extracts were analyzed with three glycan-specific methods, except for NaOH extracts that were not analyzed by high performance anionic exchange chromatography with pulsed amperometric detection (HPAEC-PAD) due to the high molarity of the solvent. See specifics of the extraction procedure and glycan analyses in Material and methods.

**Table S1.** $^{14}C$ ages of top and near to the bottommost layers of the three cores studied. Age of the top and bottom layers of the central anoxic fraction of the OAO core were determined as well.

| Core ID | Core depth (cm) | Fraction modern carbon | $^{14}C$ age (years BP) | Core layer |
|---|---|---|---|---|
| GeoB7802-1 | 1 | 0.9140 ± 0.0081 | 722 ± 71 | Anoxic core top |
| GeoB7802-1 | 30 | 0.7146 ± 0.0077 | 2,699 ± 86 | Anoxic core near to bottom |
| GeoB7807-2 | 0.2 | 1.0034 ± 0.0086 | modern | Oxic core top |
| GeoB7807-2 | 21 | 0.6634 ± 0.0072 | 3,296 ± 87 | Oxic core near to bottom |
| GeoB7830-1 | 0.2 | 0.9973 ± 0.0088 | 22 ± 71 | OAO core top |
| GeoB7830-1 | 8 | 0.8664 ± 0.0082 | 1,152 ± 76 | OAO anoxic fraction top |
| GeoB7830-1 | 19 | 0.7123 ± 0.0070 | 2,725 ± 79 | OAO anoxic fraction bottom |
| GeoB7830-1 | 23 | 0.6245 ± 0.0071 | 3,781 ± 91 | OAO core near to bottom |

BP: before present. OAO: oxic-anoxic-oxic.

**Table S2.** Description of the sectioned layers of the Shaban Deep core, GeoB7802-1. Half of the anoxic brine core (split by the longitudinal plane) was transversely sectioned into a total of 33 layers in this study. The following information, based on visual inspection, was noted down for each sectioned layer. The here stated color brown denotes dark olive brown to dark olive grey. N/A: not available.

| Depth, cm below surface | Layer size (cm) | Description | Position in the core |
|---|---|---|---|
| 1 | 1 | N/A | Top of the core |
| 2 | 1 | Laminated | |
| 2.8 | 0.8 | N/A | |
| 4 | 1.2 | Turbid | |
| 5 | 1 | Turbid plus a little bit mixed (brown and white material) | |
| 6 | 1 | N/A | |
| 7 | 1 | Laminated | |
| 8 | 1 | Mostly brown | |
| 9 | 1 | Mix of white and brown (more brown) | |
| 10 | 1 | Mostly brown | |
| 11 | 1 | Mix of white and brown | |
| 12 | 1 | Mix of white and brown | |
| 13 | 1 | Mix of white and brown | |
| 14 | 1 | Brown with two big white layers | |
| 15 | 1 | Brown | |
| 16 | 1 | Mostly brown with a bit of white | |
| 17 | 1 | Mostly brown with a single white layer | |
| 18 | 1 | Mostly brown with two thin white layers | |
| 19 | 1 | Brown | |
| 20 | 1 | Mostly brown with a thin white layer | |
| 21 | 1 | Mostly brown with a thick (approx. 2 mm) white layer | |
| 22 | 1 | Mostly brown with two very thin white layers | |
| 23 | 1 | Brown/white/turbid with a ratio 3/1/6 | |
| 23.5 | 0.5 | Only turbid | |
| 24 | 0.5 | Mostly turbid | |
| 25 | 1 | Almost completely brown | |
| 26 | 1 | Almost completely brown | |
| 27 | 1 | Brown with thin layers of white (5 thin layers) | |
| 28 | 1 | Only white (with a very thin brown layer at the edge) | |
| 29 | 1 | All brown | |
| 30 | 1 | Brown with two white layers | |
| 31 | 1 | Brown with two very thin white layers | |
| 31.5 | 0.5 | Brown | Bottom of the core |

**Table S3.** Total organic carbon (TOC) and calcium carbonate ($CaCO_3$) values of the GeoB7802-1 anoxic brine core presented as percentage per sediment dry weight. n.m.: not measured.

| Depth, cm below surface | TOC % | $CaCO_3$ % |
|---|---|---|
| 1 | 2.04 | 45.98 |
| 2 | 1.78 | 43.15 |
| 2.8 | 1.84 | 42.65 |
| 4 | 2.15 | 47.31 |
| 5 | 1.76 | 48.31 |
| 6 | 1.66 | 43.57 |
| 7 | n.m. | n.m. |
| 8 | 2.03 | 40.98 |
| 9 | 1.93 | 57.31 |
| 10 | 2.03 | 39.40 |
| 11 | 1.91 | 45.07 |
| 12 | 2.13 | 46.48 |
| 13 | 1.53 | 39.32 |
| 14 | 1.27 | 45.40 |
| 15 | 1.45 | 41.90 |
| 16 | 1.75 | 41.07 |
| 17 | 1.6 | 37.40 |
| 18 | 1.7 | 42.65 |
| 19 | 2.01 | 41.40 |
| 20 | 1.93 | 43.23 |
| 21 | 1.71 | 42.98 |
| 22 | 2.04 | 40.82 |
| 23 | 1.15 | 36.40 |
| 23.5 | 1.51 | 43.07 |
| 24 | 1.59 | 47.23 |
| 25 | 1.78 | 35.82 |
| 26 | 1.14 | 50.81 |
| 27 | 1.98 | 40.48 |
| 28 | 0.33 | 60.31 |
| 29 | 2.05 | 39.57 |
| 30 | 2.03 | 42.82 |
| 31 | 1.99 | 44.48 |
| 31.5 | 1.82 | 39.40 |

**Table S4.** Development and optimization of a polysaccharide extraction protocol for brine sediments. Overview of the parameters and conditions tested in this study. Details of the optimized extraction procedure are depicted in Material and methods.

| Parameters | Tested conditions |
| --- | --- |
| Extraction method | Sonication vs. shaking on a TissueLyser II from Qiagen (samples shaken at 30/s for 2 min and then at 6/s for 2 h, at room temperature) |
| Extraction solvents | MilliQ water, NaCl, EDTA, HCl, NaOH |
| Molarity of the solvents | NaCl: 0.5 M, 2 M, 5 M; EDTA: 0.1 M, 0.2 M, 0.5 M; HCl: 1 M; NaOH: 0.5 M, 2 M, 4 M |
| Ratio of solvent volume per sediment dry weight | The following solvent volumes (µL) were tested: 100, 200, 300, 400, 500, 600. These volumes (for MilliQ, EDTA and NaOH) were added to 10 mg of dry sediment |
| Sonication extraction time | 30 min, 60 min, 120 min |
| Alcohol-insoluble residue (AIR) | AIR prior to extraction vs. no AIR prior to extraction |

**Table S5.** Sum (addition of the three extracts per layer) of total carbohydrates (TCHO) presented as the average throughout the core. In the case of the oxic-anoxic-oxic (OAO) core the average includes the layers from each of the three fractions.

| **Core ID** | **Core name, fraction** | **TCHO Sum average (mg g dry weight$^{-1}$)** |
|---|---|---|
| GeoB7802-1 | Anoxic core | 5.83 ± 1.32 |
| GeoB7807-2 | Oxic core | 0.65 ± 0.23 |
| GeoB7830-1 | OAO core, upper oxic | 0.97 ± 0.20 |
| GeoB7830-1 | OAO core, central anoxic | 3.35 ± 0.78 |
| GeoB7830-1 | OAO core, lower oxic | 1.12 ± 0.27 |

**Table S6.** Specificities of the probes used in this study for analyzing the sediment extracts.

| Probe | Recognized glycan epitope | Reference |
|---|---|---|
| JIM5 | Partially methyl-esterified/de-esterified HG | (9) |
| JIM7 | Partially methyl-esterified HG | (9) |
| LM18 | Partially methyl-esterified/de-esterified HG | (10) |
| LM19 | Partially methyl-esterified/de-esterified HG | (10) |
| LM20 | Partially methyl-esterified HG | (10) |
| LM7 | Non-blockwise partially methyl-esterified HG | (11) |
| LM8 | Xylogalacturonan | (12) |
| 2F4 | HG cross-linked through calcium ions | (13) |
| INRA-RU1 | Rhamnogalacturonan I backbone | (14) |
| INRA-RU2 | Rhamnogalacturonan I backbone | (14) |
| LM16 | Galactosyl residue(s) on rhamnogalacturonan I | (15) |
| LM5 | (1→4)-β-D-galactan | (16) |
| LM6 | (1→5)-α-L-arabinan | (17) |
| LM21 | (1→4)-β-D-(galacto)(gluco)mannan | (18) |
| LM22 | (1→4)-β-D-(galacto)(gluco)mannan | (18) |
| BS-400-4 | (1→4)-β-D-(galacto)(gluco)mannan | (19) |
| BS-400-2 | (1→3)-β-D-glucan | (20) |
| BS-400-3 | (1→3)(1→4)-β-D-glucan | (21) |
| *CBM3a | Cellulose | (22) |
| LM15 | Xyloglucan (XXXG motif) | (23) |
| LM24 | Galactosylated xyloglucan | (24) |
| LM25 | Xyloglucan (XXXG motif, both galactosylated and non-galactosylated) | (24) |
| *LM10 | (1→4)-β-D-xylan | (25) |
| *LM11 | (1→4)-β-D-xylan/arabinoxylan | (25) |
| *INRA-AX1 | (1→4)-β-D-xylan/arabinoxylan | (26) |
| LM23 | Xylosyl residues | (24) |
| LM12 | Feruloylated polymers | (24) |
| LM2 | β-linked GlcA in arabinogalactan protein | (27) |
| LM14 | GlcA in arabinogalactan protein | (24, 28) |
| JIM13 | Arabinogalactan protein glycan | (29) |
| JIM14 | Arabinogalactan protein glycan | (29) |
| JIM16 | Arabinogalactan protein glycan | (29) |
| LM1 | Extensin | (30) |
| JIM20 | Extensin | (31) |
| BAM1 | Un-sulfated epitope present in sulfated fucan | (32) |
| BAM2 | Sulfated epitope present in sulfated fucan | (32) |
| BAM3 | Possibly sulfated epitope present in sulfated fucan | (32) |
| BAM4 | Sulfated epitope present in sulfated fucan | (32) |
| BAM6 | Alginate - mannuronate-rich epitope | (33) |
| BAM7 | Alginate - mannuronate-guluronate | (33) |
| BAM8 | Alginate - mannuronate-guluronate | (33) |
| BAM9 | Alginate - mannuronate-guluronate | (33) |
| BAM10 | Alginate - mannuronate-guluronate | (33) |
| BAM11 | Alginate - ~7 guluronate residues | (33) |

*LM10, LM11, INRA-AX1 and CBM3a were not quantified because of background noise signal. All probes used in this study are monoclonal antibodies except for CBM3a, which is a carbohydrate binding module (CBM). HG: homogalacturonan. G: glucose. X: xylose. GlcA: glucuronic acid.

#### Supplementary Information references


1. G. Pautot, P. Guennoc, A. Coutelle, N. Lyberis, Discovery of a large brine deep in the northern Red Sea. *Nature* **310**, 133–136 (1984).
2. M. Hartmann, J. C. Scholten, P. Stoffers, F. Wehner, Hydrographic structure of brine-filled deeps in the Red Sea - new results from the Shaban, Kebrit, Atlantis II, and Discovery Deep. *Mar. Geol.* **144**, 311–330 (1998).
3. J. Pätzold, G. Bohrmann, C. Hübscher, "Black Sea-Mediterranean-Red Sea, Cruise No. 52, January 2-March 27, 2002. METEOR-Berichte 03-2, Leg 3. Universität Hamburg" (2003).
4. I. A. Seeberg-Elverfeldt, C. B. Lange, H. W. Arz, J. Pätzold, J. Pike, The significance of diatoms in the formation of laminated sediments of the Shaban Deep, Northern Red Sea. *Mar. Geol.* **209**, 279–301 (2004).
5. I. A. Seeberg-Elverfeldt, C. B. Lange, J. Pätzold, G. Kuhn, Laminae type and possible mechanisms for the formation of laminated sediments in the Shaban Deep, northern Red Sea. *Ocean Sci.* **1**, 113–126 (2005).
6. S. S. Sofianos, W. E. Johns, Observations of the summer Red Sea circulation. *J. Geophys. Res. Ocean.* **112**, C06025 (2007).
7. H. W. Arz, F. Lamy, J. Pätzold, A pronounced dry event recorded around 4.2 ka in brine sediments from the northern Red Sea. *Quat. Res.* **66**, 432–441 (2006).
8. A. Ehrhardt, C. Hübscher, "The Northern Red Sea in transition from rifting to drifting - lessons learned from Ocean Deeps" in *The Red Sea*, N. M. A. Rasul, I. C. F. Stewart, Eds. (Springer Berlin Heidelberg, 2015), pp. 99–121.
9. M. H. Clausen, W. G. T. Willats, J. P. Knox, Synthetic methyl hexagalacturonate hapten inhibitors of anti-homogalacturonan monoclonal antibodies LM7, JIM5 and JIM7. *Carbohydr. Res.* **338**, 1797–1800 (2003).
10. Y. Verhertbruggen, S. E. Marcus, A. Haeger, J. J. Ordaz-Ortiz, J. P. Knox, An extended set of monoclonal antibodies to pectic homogalacturonan. *Carbohydr. Res.* **344**, 1858–1862 (2009).
11. W. G. T. Willats, *et al.*, Modulation of the degree and pattern of methyl-esterification of pectic homogalacturonan in plant cell walls: implications for pectin methyl esterase action, matrix properties, and cell adhesion. *J. Biol. Chem.* **276**, 19404–19413 (2001).
12. W. G. T. Willats, *et al.*, A xylogalacturonan epitope is specifically associated with plant cell detachment. *Planta* **218**, 673–681 (2004).
13. F. Liners, J.-J. Letesson, C. Didembourg, P. Van Cutsem, Monoclonal antibodies against pectin: recognition of a conformation induced by calcium. *Plant Physiol.* **91**, 1419–1424 (1989).
14. M.-C. Ralet, O. Tranquet, D. Poulain, A. Moïse, F. Guillon, Monoclonal antibodies to rhamnogalacturonan I backbone. *Planta* **231**, 1373–1383 (2010).
15. Y. Verhertbruggen, *et al.*, Developmental complexity of arabinan polysaccharides and their processing in plant cell walls. *Plant J.* **59**, 413–425 (2009).
16. L. Jones, G. B. Seymour, J. P. Knox, Localization of pectic galactan in tomato cell walls using a monoclonal antibody specific to (1→4)-β-D-galactan. *Plant Physiol.* **113**, 1405–1412 (1997).
17. W. G. T. Willats, S. E. Marcus, J. P. Knox, Generation of a monoclonal antibody specific to (1→5)-alpha-L-arabinan. *Carbohydr. Res.* **308**, 149–152 (1998).
18. S. E. Marcus, *et al.*, Restricted access of proteins to mannan polysaccharides in intact plant cell walls. *Plant J.* **64**, 191–203 (2010).
19. F. A. Pettolino, *et al.*, A (1→4)-β-mannan-specific monoclonal antibody and its use in

the immunocytochemical location of galactomannans. *Planta* **214**, 235–242 (2001).
20. P. J. Meikle, I. Bonig, N. J. Hoogenraad, a E. Clarke, B. a Stone, The location of (1→3)-β-glucans in the walls of pollen tubes of Nicotiana alata using a (1→3)-β-glucan-specific monoclonal antibody. *Planta* **185**, 1–8 (1991).
21. P. J. Meikle, N. J. Hoogenraad, I. Bonig, A. E. Clarke, B. A. Stone, A (1→3,1→4)-β-glucan-specific monoclonal antibody and its use in the quantitation and immunocyto-chemical location of (1→3,1→4)-β-glucans. *Plant J.* **5**, 1–9 (1994).
22. A. W. Blake, *et al.*, Understanding the biological rationale for the diversity of cellulose-directed carbohydrate-binding modules in prokaryotic enzymes. *J. Biol. Chem.* **281**, 29321–29329 (2006).
23. S. E. Marcus, *et al.*, Pectic homogalacturonan masks abundant sets of xyloglucan epitopes in plant cell walls. *BMC Plant Biol.* **8** (2008).
24. H. L. Pedersen, *et al.*, Versatile high resolution oligosaccharide microarrays for plant glycobiology and cell wall research. *J. Biol. Chem.* **287**, 39429–39438 (2012).
25. L. McCartney, S. E. Marcus, J. P. Knox, Monoclonal antibodies to plant cell wall xylans and arabinoxylans. *J. Histochem. Cytochem.* **53**, 543–546 (2005).
26. F. Guillon, *et al.*, Generation of polyclonal and monoclonal antibodies against arabinoxylans and their use for immunocytochemical location of arabinoxylans in cell walls of endosperm of wheat. *J. Cereal Sci.* **40**, 167–182 (2004).
27. E. A. Yates, *et al.*, Characterization of carbohydrate structural features recognized by anti-arabinogalactan-protein monoclonal antibodies. *Glycobiology* **6**, 131–139 (1996).
28. I. Moller, *et al.*, High-throughput screening of monoclonal antibodies against plant cell wall glycans by hierarchical clustering of their carbohydrate microarray binding profiles. *Glycoconj. J.* **25**, 37–48 (2008).
29. J. P. Knox, P. J. Linstead, J. Peart, C. Cooper, K. Roberts, Developmentally regulated epitopes of cell surface arabinogalactan proteins and their relation to root tissue pattern formation. *Plant J.* **1**, 317–326 (1991).
30. M. Smallwood, H. Martin, J. P. Knox, An epitope of rice threonine- and hydroxyproline-rich glycoprotein is common to cell wall and hydrophobic plasma-membrane glycoproteins. *Planta* **196**, 510–522 (1995).
31. M. Smallwood, *et al.*, Localization of cell wall proteins in relation to the developmental anatomy of the carrot root apex. *Plant J.* **5**, 237–246 (1994).
32. T. A. Torode, *et al.*, Monoclonal antibodies directed to fucoidan preparations from brown algae. *PLoS One* **10**, e0118366 (2015).
33. T. A. Torode, *et al.*, Dynamics of cell wall assembly during early embryogenesis in the brown alga Fucus. *J. Exp. Bot.* **67**, 6089–6100 (2016).
